# Histidine Protonation Controls Structural Heterogeneity in the Cyanobacteriochrome AnPixJg2

**DOI:** 10.1101/2020.08.21.260158

**Authors:** Aditya G. Rao, Christian Wiebeler, Saumik Sen, David S. Cerutti, Igor Schapiro

**Affiliations:** Fritz Haber Center for Molecular Dynamics Research, Institute of Chemistry, The Hebrew University of Jerusalem, Jerusalem 91904, Israel; Department of Chemistry and Chemical Biology, Rutgers University, USA

**Author notes:** Corresponding author: Dr. Igor Schapiro, Fritz Haber Center for Molecular Dynamics Research, Institute of Chemistry, The Hebrew University of Jerusalem, Jerusalem 91904, Israel.

**Keywords:** molecular dynamics, theoretical spectroscopy, red/green CBCR, QM/MM

## Abstract

Cyanobacteriochromes are compact and spectrally diverse photoreceptor proteins that bind a linear tetrapyrrole as a chromophore. They show photochromicity by having two stable states that can be interconverted by the photoisomerization of the chromophore. Hence, these photochemical properties make them an attractive target for biotechnological applications. However, their application is impeded by structural heterogeneity that reduces the yield of the photoconversion. The heterogeneity can originate either from the chromophore structure or the protein environment. Here, we study the origin of the heterogeneity in AnPixJg2, a representative member of the red/green cyanobacteriochrome family, that has a red absorbing parental state and a green absorbing photoproduct state. Using molecular dynamics simulations and umbrella sampling we have identified the protonation state of a conserved histidine residue as a trigger for structural heterogeneity. When the histidine is in a neutral form, the chromophore structure is homogenous, while in a positively charged form, the chromophore is heterogeneous with two different conformations. We have identified a correlation between the protonation of the histidine and the structural heterogeneity of the chromophore by detailed characterization of the interactions in the protein binding site. Our findings reconcile seemingly contradicting spectroscopic studies that attribute the heterogeneity to different sources. Furthermore, we predict that circular dichroism can be used as a diagnostic tool to distinguish different substates.

**Significance statement:** Cyanobacteriochromes are photoreceptor proteins that have attracted attention for their immense potential in bioimaging and optogenetics applications. This is due to their desirable properties such as compactness, photochromicity and diverse spectral tuning. Despite these advantages, nature has set a limitation in the form of structural heterogeneity that presents a drawback for its application in biotechnology. We have identified a histidine residue in the vicinity of the chromophore as the origin of the heterogeneity in red/green CBCRs. The protonation state of this conserved histidine alters an extended network of protein-chromophore interactions and induces heterogeneity. Furthermore, theoretical CD spectroscopy has revealed easy identification of heterogeneity. Hence, our study paves the way for rational design and optimization of protein properties.

## Introduction

Phytochromes constitute a large family of photoreceptor proteins found in plants, bacteria and fungi (1). These photoreceptor proteins consist of three domains: PAS (Per-ARNT-Sim), GAF (cGMP-phosphodiesterase/adenylate cyclase/FhlA) and PHY (phytochrome specific). The GAF domain binds a linear tetrapyrrole chromophore through a conserved cysteine residue (2) nevertheless all three domains are essential for the complete photochemistry(3). Upon light absorption, a photocycle is initialized by the photoisomerization of the tetrapyrrole chromophore. A typical photocycle of phytochromes exhibit a reversible conversion from a red-absorbing parental state (Pr) to a far-red absorbing photoproduct state (Pfr). The red/far-red color tuning is conserved among the phytochromes, despite their wide distribution in various organisms. A much wider spectral tunability is available in a subfamily of phytochromes recently discovered in cyanobacteria and named cyanobacteriochromes (CBCRs) (3, 4). These proteins are spectrally diverse allowing color detection from the near-UV to the near-IR range (3). This wide range of absorption covered by CBCRs makes them an attractive target for non-invasive optical imaging and optogenetics (5). In addition to the spectral properties, the CBCRs require only the GAF domain for photoreception which typically consists of less than 200 amino acids. Therefore, the compact size together with the spectral tuning makes CBCRs excellent candidates for biotechnological applications (6).

There are at least four CBCR subfamilies reported in the literature (3). One of the most studied subfamilies has a red-absorbing parental state (Pr) like the canonical phytochromes but a green-absorbing photoproduct state (Pg) (2, 5, 7–30). This strongly blue-shifted photoproduct contrasts with the far-red shifted product in canonical phytochromes which brought the unusual spectral tuning in the focus of numerous studies. Several representatives of this red/green CBCR subfamily have been extensively characterized by spectroscopic methods: AnPixJg2 (2, 17, 24–29), NpR6012g4 (7–16, 18, 30) and Slr1393g3 (5, 19–23). Also, crystallographic structures of some representative proteins in this subfamily were solved (2, 21). The first of these crystal structures was reported by Ikeuchi and coworkers (2) for the Pr state of AnPixJg2, a phototaxis regulator (PixJ) found in *Anabaena* sp. PCC 7120. AnPixJg2 uses the chromophore phycocyanobilin (PCB) that is present in a C_5_-*Z*,syn/C_10_-*Z*,syn/C_15_-*Z*,anti (*ZZZssa*) conformation as revealed in the crystal structure (Fig. 1A). The chromophore binding site has several conserved amino acids among the red/green subfamily that are involved in different types of interactions with PCB (Fig. 1A). Some of these interactions are altered upon light-activated isomerization that takes place around the C_15_=C_16_ double bond (Fig. 1B). (17)

**Fig. 1.**
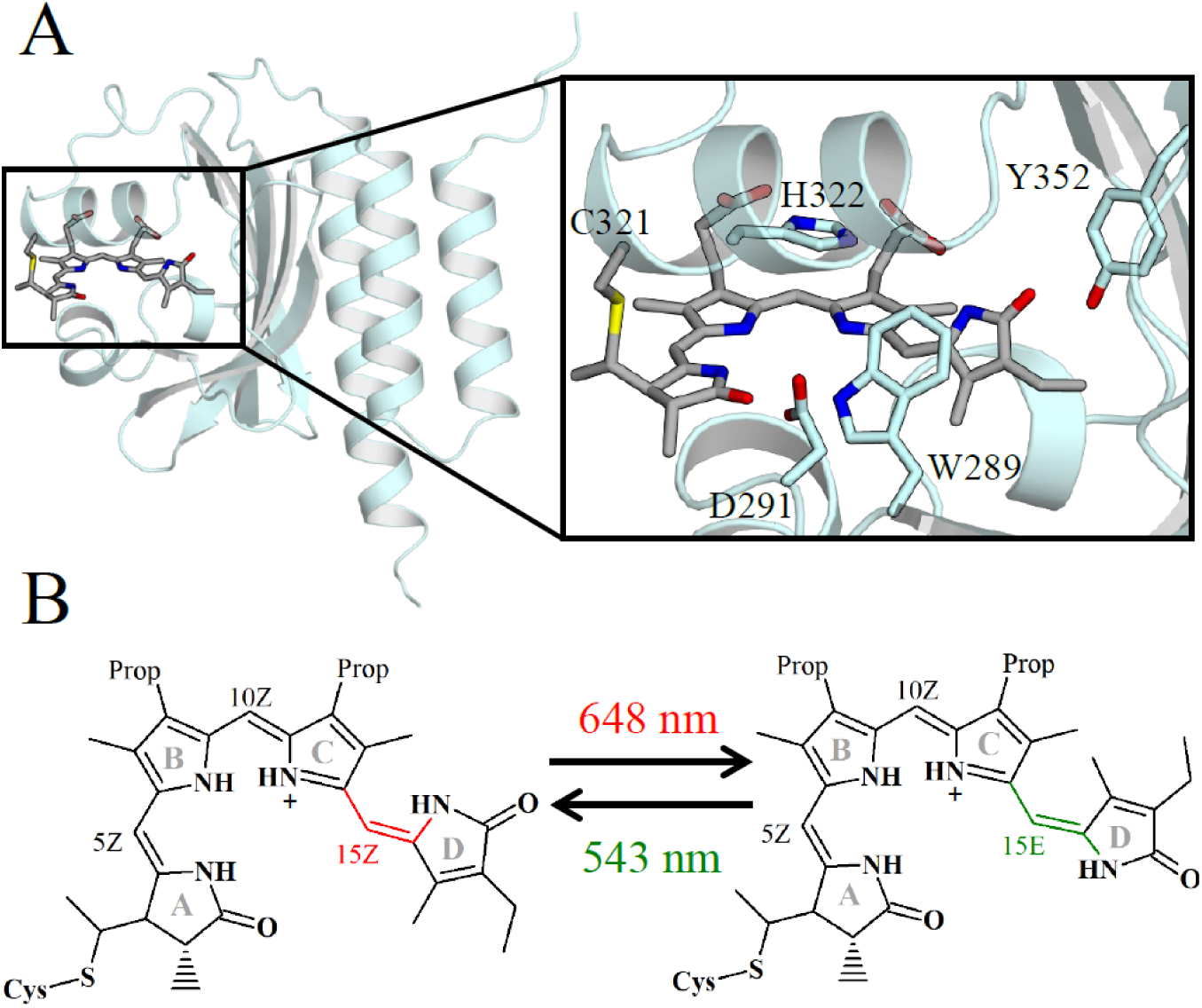
The red/green CBCR AnPixJg2. (A) Crystal structure of the Pr form of AnPixJg2 (PDB ID: 3W2Z) with the chromophore binding pocket and the critical residues (C321, H322, W289, D291 and Y352). (B) Reversible photoisomerization between the red-absorbing dark state (Pr) and the green-absorbing photoproduct (Pg).

This process triggers the photocycle that eventually yields the green-absorbing Pg state. However, despite the identical absorption maxima in the photostationary states of the red/green CBCR subfamily the conversion process and the photocycle kinetics can vary significantly. Such differences are attributed to heterogeneity in both structural and photochemical properties (14). For example, in the phylogenetically related NpR6012g4 the quantum yield of the Pr to Pg conversion is reported to be 40% (9), while in Slr1393g3 it is found to be reduced to 8% (5, 19). Since a low quantum yield can impede the use of CBCRs for biotechnological applications, the heterogeneity in CBCRs calls for a detailed investigation at the molecular level.

The structural heterogeneity can be ascribed to the presence of different chromophore conformations and different rotamers or protonation states of the amino acids in the binding site (9, 10, 14, 18, 19, 25, 29). In this context the orientation of the D-ring with respect to the plane formed by the B-C rings has been referred to as the source of chromophore heterogeneity. In one substate the D-ring is above the plane formed by the central B-C rings, while in the other it is below. Rockwell et al. (31) have defined these orientations as α-facial and β-facial orientations, respectively (Fig. 2A). We adapt their notation and refer to the α-facial and β-facial orientation of the D-ring as D-α_f_ and D-β_f_, respectively. Resonance Raman studies on the Pr and Pg states of AnPixJg2 have revealed a heterogeneous chromophore only in the Pg form which has been attributed to the conformers of the C-D methine bridge (25). In contrast, an NMR study demonstrated that the chromophore is heterogeneous in both forms of AnPixJg2 (29). This finding was substantiated by classical molecular dynamics (MD) simulations of the Pr state (29) that also revealed two substates with distinct rotation of the D ring. Interestingly, in a related red/green CBCR NpR6012g4 a recent NMR study found a homogeneous chromophore and assigned the heterogeneity to different rotamers of the binding site residues(14). The presence of different residue conformers was then connected to the photochemical heterogeneity(9, 10) that is reflected in several populations that evolve on different time scales. This difference between AnPixJg2 and NpR6012g4 is surprising because these proteins share ∼60% sequence identity and ∼74% similarity as calculated using the EMBOSS-NEEDLE program (32). Their sequence alignment (Fig. S3) indicates that all key residues within the chromophore binding site (C321, H322, W289, D291 and Y352) are conserved Therefore, in the present study, we address the heterogeneity of AnPixJg2 by MD simulations and theoretical optical spectroscopy with the aim to rationalize the reported differences.

**Fig. 2.**
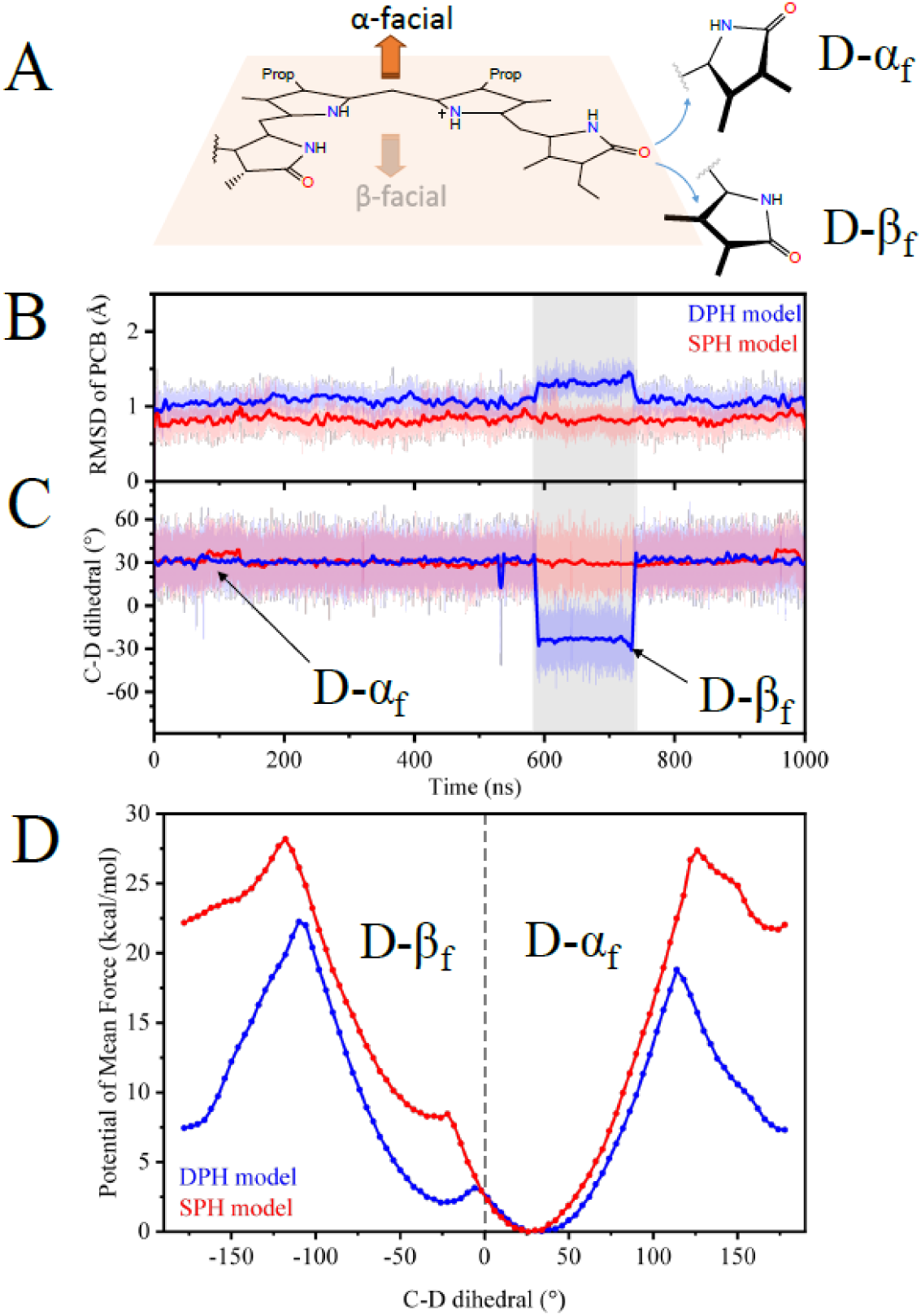
Structural flexibility of the chromophore in AnPixJg2. (A) α- and β-facial orientations of the chromophore with reference to the plane defined by the B-C rings. (B) The all-atom RMSD of PCB chromophore. (C) Dihedral angle [C_13_−C_14_−C_15_=C_16_] between C and D rings along the trajectory. (D) Potential of Mean force (PMF) obtained for SPH and DPH models. In both models, the D-β_f_ substate is relatively higher in energy compared to the D-α_f_ substate.

The starting point of our investigation is the protonation state of the critical histidine residue that was reported to play a key role in the heterogeneity of the phytochrome Cph1 (33) and lead to different hydrogen bonding networks within its binding site (34). Along similar lines, a recent computational study (35) that investigated the effect of different protonation states of the critical histidine residue in the phytochrome *Deinococcus radiodurans*, reported significant differences in the resulting non-covalent interactions between the chromophore and the protein. Hence, we have specifically studied the effect of the analog histidine in AnPixJg2 (H322) protonation on the structural dynamics of PCB and found a connection between its protonation state and the ground state heterogeneity. To this end we have derived parameters for PCB consistent with the AMBER ff14SB (36) force field, which allowed us to perform MD simulations. To characterize the protein-chromophore interactions, we have decomposed the non-covalent interactions using Symmetry Adapted Perturbation Theory (SAPT). We have also calculated UV/Vis and CD spectra using the hybrid quantum mechanics/molecular mechanics (QM/MM) method to see how our findings can be validated experimentally. The calculations were performed along the lines of our previous study for Slr1393g3 (23), i.e. geometries were sampled using molecular dynamics simulations and then excitation energies were computed using simplified time-dependent density functional theory (sTD-DFT) (37–39) and the algebraic diagrammatic construction method to second order (ADC(2)) (40).

This paper is organized as follows: First, we describe the dynamics of the chromophore in AnPixJg2 using classical MD and umbrella sampling simulations. We have carried out these simulations with H322 in two different protonation states: a positively charged histidine that is protonated at both the ε-n and δ-positions (DPH model) and a neutral histidine protonated only at the δ-position (SPH model). Second, we compare the hydrogen bonding interactions within the chromophore binding site for both models. Further, we analyze the dynamics of W289 in both models, as this “lid Trp” is ascribed a critical role in red/green CBCRs (25, 30). In the last section, we determine the spectroscopic properties for the two models and suggest the use of CD spectroscopy as a diagnostic tool to determine heterogeneity in red/green CBCRs.

## Results and Discussion

### Structural flexibility of the chromophore

To study the effect of the H322 protonation state on the conformation of AnPixJg2 in the Pr state classical MD simulations were performed. For each protonation state a 1 µs trajectory was calculated. The resulting all-atom root mean square deviation (RMSD) of the PCB relative to the starting structure is shown in Fig. 2B. In the SPH model the RMSD fluctuates around 0.8 Å, while in the DPH model two distinct values are observed. The major population fluctuates around 1.1 Å. It is associated with the α-facial orientation of the D ring (D-α_f_) that becomes apparent from the evolution of the C_13_−C_14_− C_15_=C_16_ dihedral (henceforth simply termed C-D dihedral). A minor population with an increased RMSD of 1.5 Å is found between 590 to 720 ns (Fig. 2B) and is linked to the β-facial orientation (D-β_f_). The mean value of the characteristic C-D dihedral in the D-α_f_ and D-β_f_ substates in the DPH model is + 31° and − 24°, respectively (Table S1). The presence of two substates in the DPH model is in agreement with the findings of Scarbath-Evers et al. (29). However, in our study the D-β_f_ substate was observed for ∼130 ns compared to the longer period of ∼570 ns in the aforementioned study. In contrast to the DPH model, the SPH model shows a uniform orientation of the D ring with a mean C-D dihedral of + 31°.The difference in the lifetime of the two substates for the DPH model and the lack of a second substate in the SPH model raises a question if two trajectories of 1 µs are sufficient to explore the full conformational space. To address this concern, we have performed umbrella sampling simulations.

### Umbrella sampling

For systematic exploration of the potential energy surface of the PCB chromophore we have used the umbrella sampling technique. In umbrella sampling, a selected reaction coordinate is restrained to a target value by a biasing potential. Multiple simulations are then run by scanning this reaction coordinate to generate umbrella surfaces. Using the statistics from these biased simulations, an unbiased free energy profile known as the potential of mean force (PMF) is built along the reaction coordinate with the help of the Weighted Histogram Analysis method (WHAM) (41, 42).

The PMF from umbrella sampling of SPH and DPH models is plotted along the C-D dihedral in Fig. 2D, which was chosen as a reaction coordinate. The PMF exhibits two minima in the DPH model located at + 26° and − 26°. They correspond to substates D-α_f_ and D-β_f_ already observed in the MD simulations. Substate D-α_f_ is 2 kcal.mol^-1^ lower in energy relative to substate D-β_f_. However, in the SPH model D-α_f_ is 8 kcal.mol^-1^ lower in energy than substate D-β_f_. Further, there is a negligible barrier for substate D-β_f_ such that it will instantly transit to substate D-α_f_, making it effectively the only minimum. Hence, our finding from the MD simulations is confirmed: the chromophore is homogenous with respect to the D-ring orientation in the SPH model, while it is found to be heterogenous in the DPH model. In the following we attempt to rationalize this difference at the molecular level.

The higher PMF of substate D-β_f_ in the SPH model originates in the altered interaction between the propionate of the B ring and H322. In the DPH model the positively charged H322 forms a salt bridge with carboxylate of the propionate. However, due to the neutral charge of H322 in the SPH model the propionate reorients and becomes solvent exposed (Fig. S5). In the DPH model this solvent exposed conformation is only observed at highly twisted C-D dihedral (−130° and −170°). Hence, the deprotonation of the H322 is one of the key factors in controlling the heterogeneity of the PCB chromophore in the ground state.

### Hydrogen bonding networks

We have characterized the extensive hydrogen-bonding interactions to establish a link between the chromophore structure and the flexibility of the protein binding site. For this purpose, we have calculated occurrences of hydrogen bonds for residues within the chromophore binding site (Table S2). The D-ring is mainly involved in two H-bonding interactions with (i) a water molecule forming a bridge to H322 (henceforth called “WAT1”), and (ii) hydroxyl-group of Y352.

The first interaction is analyzed using the SAPT method which allows to understand non-covalent interactions by partitioning them into various contributions. SAPT analysis requires a system of two molecules which is why we have compared the interactions between PCB and WAT1 (denoted as PCB-WAT1) as well as H322 and WAT1 (denoted as H322-WAT1) separately. These two interactions are different in the DPH (Fig 3A) and SPH (Fig. 3B) models. In the DPH model the H322-WAT1 interaction (− 10.33 kcal.mol^-1^) is stronger than PCB-WAT1 (− 4.54 kcal.mol^-1^) (Fig. S6A), while in SPH they are close-to-balance but reversed (− 4.74 kcal.mol^-1^ and − 6.88 kcal.mol^-1^, respectively) (Fig. S6C). In case of the H322-WAT1 interaction the difference between DPH and SPH models originates in the dominating electrostatic component due to the positive charge of the histidine. However, in case of the PCB-WAT1 interaction the small difference can be rationalized using the SAPT decomposition of interaction energy. The contributions from electrostatics and induction are smaller in the DPH model and cannot be compensated by the favorable reduction of the exchange interaction (Fig. S6B and S6D). Thus, the PCB-WAT1 interaction is stronger than H322-WAT1 in the SPH model that stabilizes the substate D-α_f_.

**Fig. 3.**
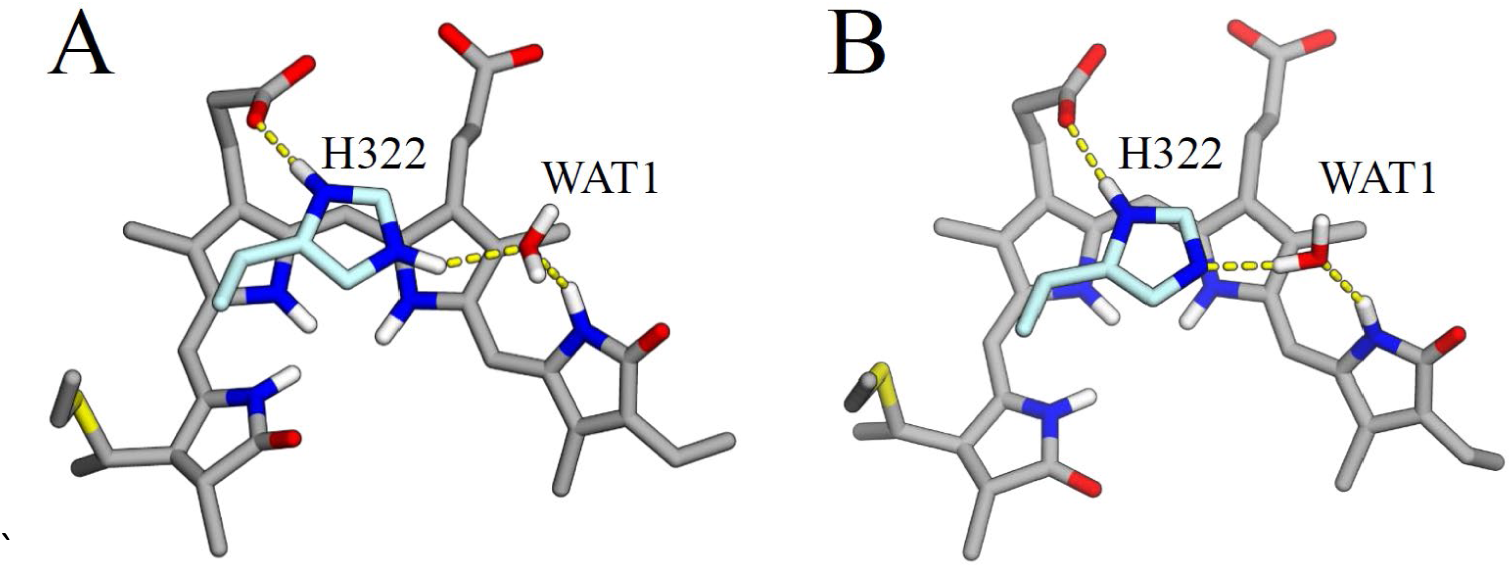
Orientation of the water molecule “WAT1” for two different protonation states of the H322. (A) In the DPH model, the oxygen atom of WAT1 has two potential H-bond donors namely ε-nitrogen of H322 and -NH moiety of the D-ring. (B) In the SPH model, the oxygen atom of WAT1 has only the -NH moiety of the D-ring as the H-bond donor. Interaction energies were calculated between PCB-WAT1 and H322-WAT1 as described in the text.

In the second interaction (ii) the occurrence of hydrogen bonding interactions between Y352 and the D-ring in the SPH model is higher (0.83) than in the DPH model (0.56 in substate D-α_f_) (Table S2). The evolution of distance between the carbonyl oxygen atom of the D-ring to the oxygen atom of the Y352 residue for DPH and SPH models is shown in Fig. 4B. In the DPH model, this interaction is present only in substate D-α_f_ and absent in substate D-β_f_. In substate D-α_f_, we observe on occasional insertion of a water molecule between Y352 and the D-ring during the simulation leading to an increase in the distance. However, this hydrogen bond is disrupted when the D-ring orients β-facially and forms a new H-bond with Y302. At the same time, the H-bonding between the propionate chain of the C-ring and Y302 is broken (Fig. 4A). When the D-ring rotates back to form substate D-α_f_, Y302 moves again closer to the propionate that is connected to ring C re-establishing the H-bonding between them (Fig 4B). In the SPH model the distances between the D-ring and the Y352 and Y302 remain unaltered.

**Fig. 4.**
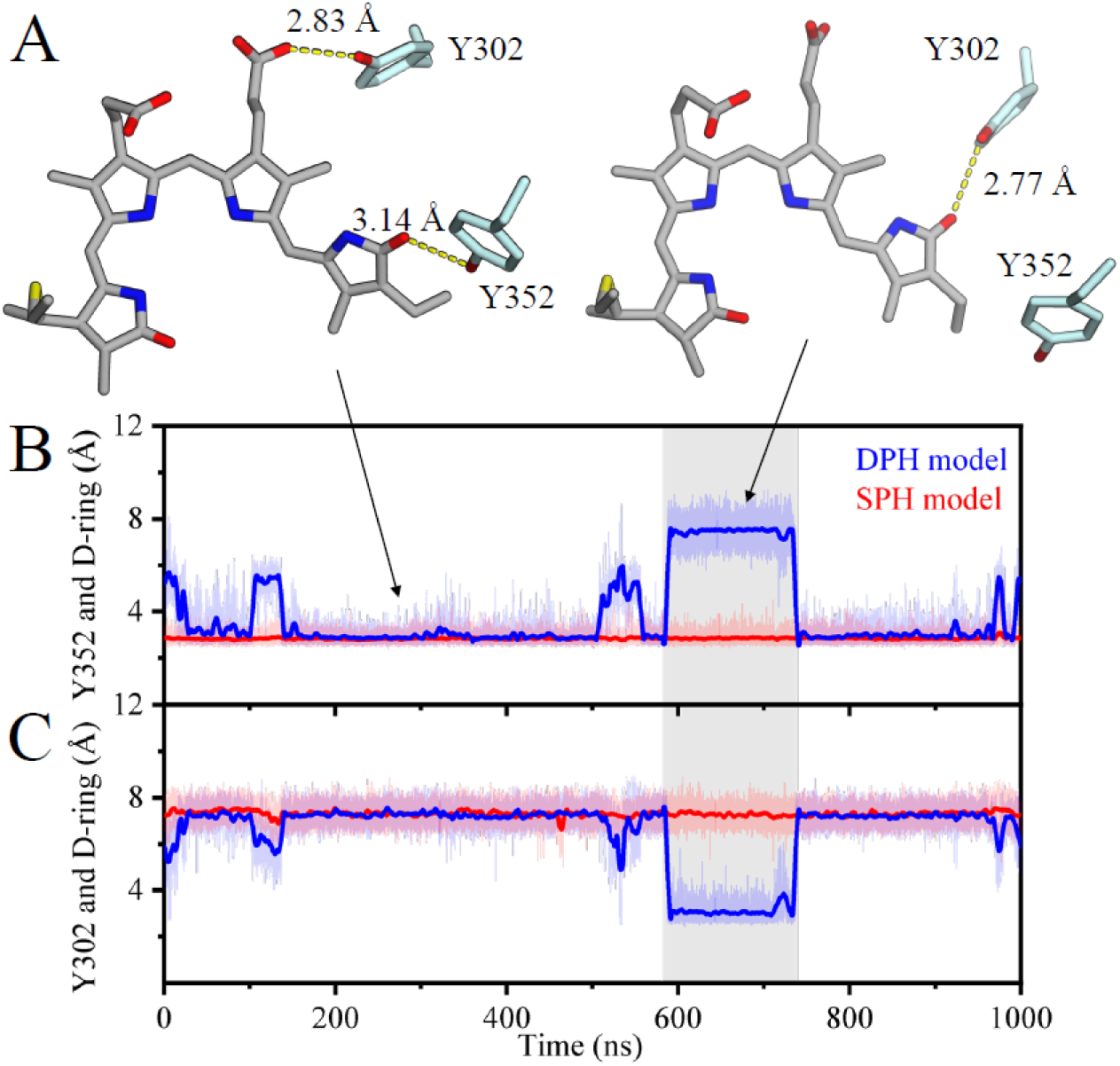
Distance between the D-ring of PCB and two tyrosines (Y302 and Y352) for different protonation states of H322. (A) In the substate D-α_f_ of the DPH model, Y302 is H-bonded to the propionate chain of the C-ring and Y352 is H-bonded to the D-ring (left). Transition to the substate D-β_f_ leads to the disruption of the H-bond with the propionate chain thereby allowing Y302 to H-bond with the D-ring and causing a movement of Y352 away from the D-ring (right). (B) Distance between the tyrosine residue Y352 and the D-ring. (C) Distance between the tyrosine residue Y302 and the D-ring.

### Tryptophan (W289) dynamics

The residue W289 is oriented parallel to the D-ring of the chromophore in the crystal structure. However, during the MD simulations of both models, the sidechain of this residue is adopting multiple conformations (Fig. 5). In the DPH model the sidechain of the W289 residue was found to switch from a parallel to a perpendicular orientation (Fig. 5D). In the perpendicular arrangement the indole group is pointing away from the A-ring. The presence of several W289 orientations is also observed in the MD simulations by Scarbath-Evers et al. (29). In the SPH model, on the contrary, W289 is found only in the parallel orientation to the D-ring of PCB. However, the hydrogen bonding between W289 and the A-ring is interrupted and then bridged by water molecules (Fig 5A). in summary, our calculations show distinct conformational dynamics of W289 for different protonation states of H322.

**Fig. 5.**
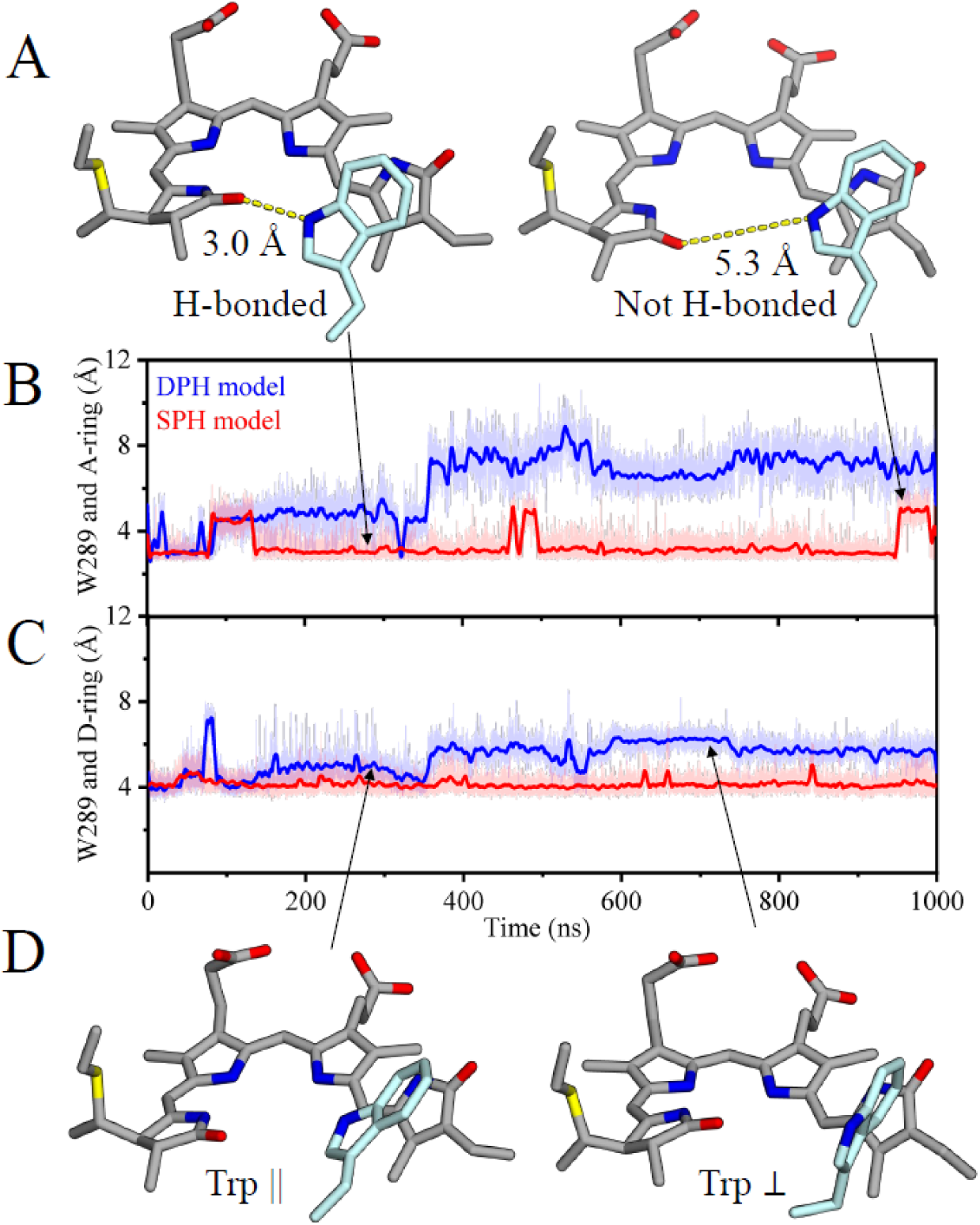
Distance between the A- and D-ring of PCB and tryptophan W289 for different protonation states of H322. (A) Snapshots of W289 (in cyan) with (left) and without (right) an H-bond to the A-ring in the SPH model. (B) Distance between W289 and the A-ring and (C) Distance between W289 and the D-ring. (D) W289 in π-stacking (left) and a perpendicular (right) orientation in DPH model.

The non-covalent interactions between PCB and W289 (denoted as PCB-W289) were analyzed using SAPT. The energy decomposition reveals a substantial contribution from the electrostatic and dispersion energies to the stabilization of the parallel conformation (Fig. S7B). The stabilizing influence of electrostatics in addition to dispersion energy in the π-stacked complexes has been observed in gas-phase studies (43, 44). A parallel conformation of W289 in the DPH model results in a mean SAPT0 energy of − 5.75 kcal.mol^-1^, while the corresponding value in the SPH model is − 10.95 kcal.mol^-1^ (Fig. S7A). The parallel conformation of the SPH model is nearly twice as stable as the DPH counterpart. The mean SAPT0 energy of the perpendicular conformation of W289 in the DPH model is − 2.15 kcal.mol^-1^. And it is only in this particular W289 orientation where we observe the substate D-β_f_. Hence, the reduced π-stacking interactions between W289 and D-ring facilitates the flip of the D-ring. This decrease in the interaction energy in the DPH model is compensated by an increase in the solvation energy of the chromophore. This is evident from the hydration pattern (water network) shown in Fig. S8. In the DPH model, a bridging water molecule between WAT1 and the central water molecule (henceforth called “WAT2”) is observed. The latter is absent in the crystal structure, that we attribute to its high mobility on the basis of our analysis. It is observed as a H-bonding partner to both the A-ring and W289 (29) with a high occurrence in the DPH model (0.79 in substate D-α_f_ and 0.72 in substate D-β_f_) and a low occurrence in the SPH model (0.06) (Table S2).

We can conclude that the W289 plays an important role in the stabilization of substate D-α_f_. However, the stability of the W289 and D-ring interaction is influenced by the protonation state of H322 and the associated hydrogen bonding network. If this residue is mutated, the missing π-stacking could alter the ratio between the two substates and therefore the structural heterogeneity.

### QM/MM Calculation of UV/Vis and CD Spectra

We have computed UV/Vis and CD spectra to probe the substates for unique spectroscopic signatures that can be used to distinguish them. To this end we have used a protocol to calculate the spectra along the lines of our recent work (23, 45, 46). In brief, to sample the geometry of PCB we have produced hybrid QM/MM trajectories with the chromophore and a few key residues in the QM region (see SI section QM/MM Molecular Dynamics). A QM description was chosen because it provides a better geometry for calculation of spectroscopic parameters than a classical force field. A 1 ns QM/MM trajectory was calculated for the SPH model and for each of the two substates of the DPH model saving snapshots every 10 ps.

The absorption spectra computed at the ADC(2)/cc-pVDZ level of theory for a QM region consisting of 66 atoms (QM66) for the DPH and SPH models are shown in Fig. 6. The absorption maxima of the lowest energy band (so-called Q-band) of substate D-α_f_ and substate D-β_f_ are found at 591 nm (2.10 eV) and 608 nm (2.04 eV), respectively. Hence, the difference in absorption between the two substates of the DPH model is small. For the SPH model the calculated absorption maximum for QM66 is 616 nm (2.01 eV). All three results are close to the experimental value of 648 nm (1.91 eV). However, the differences between the DPH and SPH models are below the accuracy of the ADC(2) method (47, 48). Extending the QM region to 106 atoms (DPH model) or 105 atoms (SPH model) did not introduce a significant shift between the absorption maxima of different substates (Fig. S9). Therefore, it is not possible to distinguish between them based on UV/Vis absorption spectra.

**Fig. 6.**
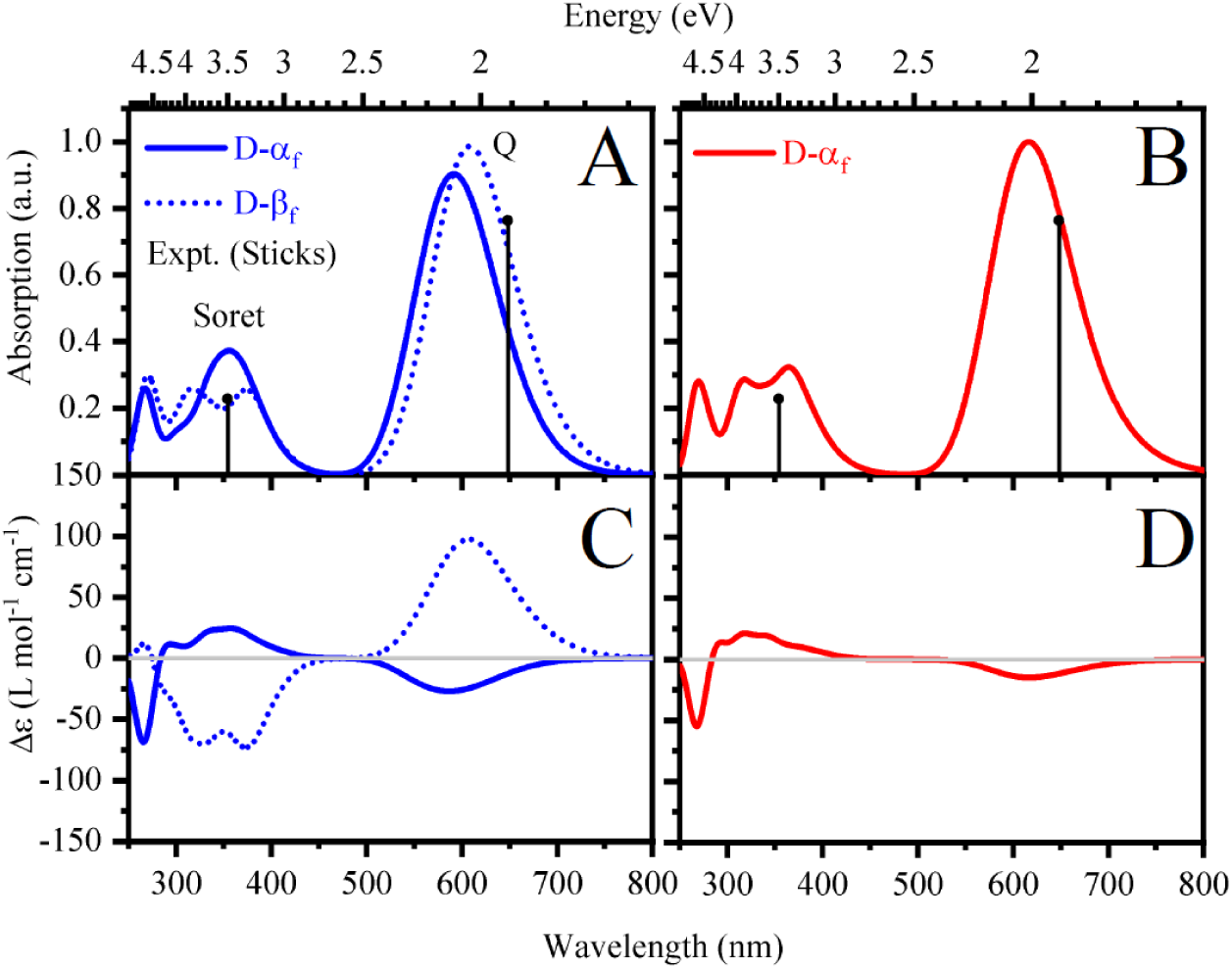
Absorption and CD spectra computed using RI-ADC(2). (A) Absorption spectra for substate D-α_f_ (solid) and substate D-β_f_ (dotted) of the DPH model (B) Absorption spectra for substate D-α_f_ (solid) of the SPH model. (C) CD spectra for substate D-α_f_ (solid) and substate D-β_f_ (dotted) of the DPH model. (D) CD spectra for substate D-α_f_ (solid) of the SPH model. The spectra are averages over 100 snapshots for each substate.

However, Rockwell et al. (31, 49) have reported a correlation between the CD rotatory strength and the orientation of the D-ring. They have shown that the sign of the Q- and Soret bands in a CD-spectrum is determined by the facial disposition of the D-ring with respect to the central, coplanar pyrrole rings B and C. Using calculations of PCB in the gas phase they have shown an inversion of the CD spectrum for different D-ring orientations (31, 50). Indeed, our calculated QM/MM CD spectra show a similar trend for the two substates in the DPH model (Fig. 6). To quantify the dependence of the rotatory strength on the peripheral rings A and D we analyzed the correlation between the excitation energy and the dihedral angle for 100 snapshots of each of the two substates (Fig. S10). In case of the Q-band the A ring has a linear correlation of 0.00, while for the D ring its − 0.91. For the Soret band the correlation coefficients are 0.07 and 0.85, respectively. Substate D-α_f_ displays a negative Q-band and a positive Soret band, while the signs are inverted for substate D-β_f_ which exhibits a much higher rotatory strength (Fig. 6C). Further, we note a three to fourfold increase in the rotatory strength for D-β_f_ compared to D-α_f_ substate. In addition, there is a relative change in the rotatory strength between the two major bands. In the DPH model, the ratio between the Q- and Soret bands is ca. 1:1 in substate D-α_f_ and 4:3 in substate D-β_f_. In the SPH model the D-α_f_ orientation displays a slightly reduced rotatory strength of both bands compared to the corresponding substate in the DPH model. More importantly, the relative rotatory strength between the Q- and Soret bands has changed from 1:1 in DPH to 2:3 in SPH model.

We compare the calculated CD spectra to the available experimental counterparts for two members of the red/green-absorbing CBCR subfamily, NpR6012g4 and Slr1393g3. The CD spectrum of NpR6012g4 (9) shows a negative Q- and a positive Soret band with a 1:2 ratio. This spectrum closely resembles the one of the SPH model that we have found to be exclusively in the D-α_f_ conformation. This finding is in line with recent NMR studies on NpR6012g4 that assign the histidine 688 (analog of H322 in AnPixJg2) to be neutral and the structure of the chromophore to be homogeneous in the ground state in agreement with our SPH model (7, 14). In contrast, AnPixJg2 was found to be heterogeneous in the ground state which agrees with the DPH model (29). Although, an experimental CD spectrum of AnPixJg2 is not available, we refer to the CD spectrum of the homolog Slr1393g3 to make a comparison for the DPH model. In the CD spectrum of Slr1393g3 (20), the Q- and Soret bands have positive rotatory strength, with a strongly diminished Q band close to zero. Rockwell et al. have systematically studied all combinations of the A and D ring disposition with coplanar B and C rings and showed that these two bands have always opposite signs. Hence, the only explanation of these two positive bands is the presence of heterogeneity in form of different D-α_f_ and D-β_f_ populations. Based on our QM/MM simulations of the DPH model we predict the ratio of the two bands in the CD spectrum of AnPixJg2 to be significantly altered as compared to NpR6012g4 and to be more similar to Slr1393g3. The absolute proportion of the Q- and Soret bands will ultimately depend on the relative population of the two substates.

## Conclusions

Hence, our calculations point towards a key role of the histidine in structural heterogeneity of red/green CBCRs. In a singly protonated form, the binding site is homogeneous with respect to the PCB conformation, i.e. only one minimum is found in the umbrella sampling calculation. However, in case of a double protonated histidine specific protein interactions result in two distinct substates. We find that CD spectroscopy can serve as a diagnostic to distinguish different substates by the sign of the CD peaks and it can also differentiate histidine protonation by the relative ratio between the Q- and Soret bands. The understanding of the molecular origin of the heterogeneity allows to rationalize recent experimental studies on two representatives of red/green CBCR subfamily. Despite having only one amino acid difference in the chromophore binding site NpR6012g4 represents an SPH model, while AnPixJg2 is an example of the DPH. This is corroborated by comparison to experimental NMR and CD spectra.

## Methodology

The detailed explanation of the parameterization of the PCB chromophore, protocols of the molecular dynamics and spectroscopy can be found in the Supporting Information (SI).

## Acknowledgments

This project has received funding from the European Research Council (ERC) under the European Union’s Horizon 2020 research and innovation programme (grant agreement No 678169, ERC Starting Grant “PhotoMutant”). C.W. is grateful for funding by the German Research Foundation (DFG) (reference numbers: WI 4853/1-1 and WI 4853/2-1). He also acknowledges computing resources provided by the Paderborn Center for Parallel Computing (PC2).

