## Supporting Information for "Histidine Protonation Controls Structural Heterogeneity in the Cyanobacteriochrome AnPixJg2"

Dr. Igor Schapiro

**This PDF file includes:**

Supplementary text
Figs. S1 to S10
Tables S1 to S5
References for SI reference citations

**Table of Contents:**


| | | |
|---|---|---|
| 1. | Parameterization of Phycocyanobilin | 3 - 5 |
| 2. | Computational Methodology | 6 - 8 |
| 3. | Supplementary Figures | 9 - 18 |
| 4. | Supplementary Tables | 19 - 24 |
| 5. | References | 25 - 26 |

### 1. Parametrization of Phycocyanobilin

In order to sample the conformation of the AnPixJg2 protein on a long-time scale using a protein force field, a set of parameters is required for phycocyanobilin (PCB). Previously, PCB parameters were reported for the CHARMM22 force field (1) and a preliminary set compatible with AMBER ff99SB (2) have been obtained by Kabasakal et al.(3). However, we have derived such parameters compatible with the AMBER ff14SB force field using the mdgx (4, 5) tool of the AMBER16 program suite.

*Definition of the AMBER ff14SB potential energy function*

The AMBER ff14SB potential energy function consists of the following bonded and non-bonded terms:

$$\boldsymbol{E_B} = \frac{1}{2} k_B (r - r_0)^2 \tag{1}$$

$$\boldsymbol{E_w}\ (\boldsymbol{\theta}) = \frac{1}{2} k_w (\theta - \theta_0)^2 \tag{2}$$

$$\boldsymbol{E_T}\ (\boldsymbol{\omega}) = \sum_j \frac{1}{2} k_j \big(1 - \cos(j\omega - \gamma)\big) \tag{3}$$

$$\boldsymbol{E_{el}}\ (\boldsymbol{r_{AB}}) = \frac{Q_A Q_B}{\varepsilon R_{AB}} \tag{4}$$

$$\boldsymbol{E_{vdW}}\ (\boldsymbol{R_{AB}}) = \left(\frac{a_{AB}}{R_{AB}^{12}} - \frac{b_{AB}}{R_{AB}^{6}}\right) \tag{5}$$

The energy of bonded interactions is a sum of energies of the bond terms ($E_B$), angle terms ($E_w$) and the dihedral angle terms ($E_T$) whereas the energy of the non-bonded interactions are a sum of the energies of electrostatic ($E_{El}$) and van-der-Waals interactions ($E_{vdW}$). The $k_B$, $k_w$ and $k_j$ denote the force constants for bonds, angles and dihedrals. $r$, $\theta$ and $\omega$ are the bond distances, bond angles and dihedral angles, respectively. $r_0$ and $\theta_0$ represent the corresponding optimized values. $j$ is the dihedral periodicity and $\gamma$ is the phase angle. The electrostatic interactions depend on the individual atomic charges ($Q_A$ and $Q_B$) separated by a distance $R_{AB}$ and the dielectric constant $\varepsilon$. The van-der-Waals interactions are described by a Lennard-Jones potential consisting of the terms $a_{AB}$ and $b_{AB}$.

*Fragmentation of Phycocyanobilin*

The crystal structure of the $P_r$ form of AnPixJg2 (PDB ID: 3W2Z) contains the PCB chromophore in a *ZZZssa* conformation. A geometry optimization of this conformation in the gas phase resulted a highly distorted structure, which originates in the attraction between the negatively charged propionic side chains and the positively charged pyrrole rings. Such intramolecular interactions occur in the gas phase because omitting the protein environment removes the partners of the charged functional groups of the chromophore. Hence, we adapted the scheme of chromophore fragmentation introduced by Mroginski et al. (1), shown in Fig. S1. To this end the propionate chains were truncated and the bonds were capped by methyl groups, resulting in fragment P1 in Fig. S1. To evaluate the angle and dihedral terms of the methine bridges, the PCB chromophore was further divided into 3 fragments (Fig. S1). The resulting parameters were then transferred to the full chromophore.

Each of the fragment was composed of two rings: fragment P2 consists of rings A and B, P3 consists of B and C rings and P4 of rings C and D. GAFF atom types were assigned to all atoms except the atoms of the cysteine moiety where AMBER atom types were used. To decouple parameters from different torsions having same atom types, four new atom types namely m1, m2, m3 and m4 were introduced. A list of atoms, their corresponding atom types and partial charges are given in Table S3. Antechamber was used to derive an initial set of charges and parameters for the fragments using GAFF2 (6) and ff14SB force field. These initial parameters were refined using the mdgx tool of the AMBER16 program.

*Target Data*

The partial charges were derived at the HF/6-31G* level of theory for all four fragments and derived charges for the larger fragment (P1) were later transferred to the full chromophore. After the charge derivation, the angle and dihedral parameters were derived at the MP2/cc-pVTZ level of theory from MM-minimized conformations of only the smaller fragments P2, P3 and P4.

*Derivation of Partial Charges*

Using the initial guess from Antechamber, multiple conformations of each fragment were generated. This is necessary to ensure that partial charges are not dependent on a single conformation. Vacuum charge densities on an electrostatic potential grid for each conformation were generated. Using the RESP method, these densities were fitted in a least squared manner to reproduce the electrostatic potential of the molecule. All four fragments were taken together during the fitting process. The partial charges of the propionate chains were directly taken from aspartic acid of the ff14SB force field, as the side chains were not included in our models.

*Derivation of Bonded Parameters*

Multiple conformations of the fragments (P2, P3 and P4) were generated by applying restraints on the dihedrals followed by energy-minimization. Several conformations of the fragments were generated with restraints on different dihedrals to ensure effective sampling of the various degrees of freedom in the chromophore. MP2 energies were then calculated using the force field optimized conformations so that bonded terms of the AMBER potential energy function could be fitted to the resulting MP2 energies. Iterations of the above process were done to cover the full conformational space of the chromophore and to obtain a good fit between the QM and MM data. In contrast, the parameters for the propionate chains were taken from the aspartic acid residue in the GAFF2 force field. The angles and dihedrals that were parameterized are listed in Table S4 and Table S5 respectively.

### 2. Computational Methodology

*Classical Molecular Dynamics simulations*

The starting geometry of the protein was based on the red absorbing form ($P_r$) of the AnPixJg2 crystal structure (PDB ID: 3W2Z). Missing hydrogen atoms were added to the crystal structure using the *tleap* program of AMBER 16. The protonation states of all titratable residues were considered at the pH 7.0. H322 was modelled in two different protonation states: doubly (δ- and ε- position) and singly (δ- position only) protonated, while H318 was modelled only in the δ- protonated state. The model with the doubly protonated H322 was named "DPH model" and the model with the singly protonated H322 "SPH model". The propionate chains attached to the B and C rings of PCB were modelled in the deprotonated form. The protein was solvated in a rectangular box of TIP3P water molecules using the *solvatebox* plugin of *tleap*, with a distance of at least 10 Å between the atoms and the boundaries of the box. The *addions2* plugin of *tleap* was used to neutralize the system by adding Na+ ions. In all classical MD simulations, the bonds involving the hydrogen atoms were constrained using the SHAKE algorithm allowing a time step of 2 fs. The cut-off for non-bonded interactions was 8.0 Å and long-range interactions were treated using the Particle Mesh Ewald algorithm. A Langevin thermostat with a collision frequency of 1 $ps^{-1}$ was used for temperature control in all simulations. VMD (Visual Molecular Dynamics) (7) was used for visualization of the trajectories from the simulation. PCB was described using the aforementioned derived parameters and the simulations were carried out using the AMBER 16 program.

At the beginning of the simulation**,** the solvent was minimized in 10000 steps with restraints of 100 kcal/mol $Å^2$ on all atoms of protein and PCB. The full system was then gradually heated from 100 K to 300 K in 1 ns with restraints on protein and PCB in NVT conditions. The density of the solvent was then gradually equilibrated for 5 ns in NPT conditions. The equilibration was extended for another 5 ns with weaker restraints of 10 kcal/mol $Å^2$. The system was then optimized for 10000 steps with restraints only on the protein backbone. Further, MD of 5 ns each was carried out with weakened restraints of 10 kcal/mol $Å^2$, 1 kcal/mol $Å^2$ and 0.1 kcal/mol $Å^2$ on the backbone. Before moving to the final production run of 1 μs, an unrestrained MD of 50 ns was carried out. The residues

P215-R231 and Q364-T388 were discarded from all structural analyses since we are simulating a monomeric form of AnPixJg2.

*Umbrella Sampling simulations*

The reaction coordinate in our simulations was the dihedral along the single bond between the C and D rings. A harmonic restraint of 100 kcal/mol rad$^2$ was applied to the dihedral. The dihedral was then incremented from -180° to +180° in steps of 10° resulting in a total of 37 windows. Each window was subjected to heating for 1 ns, equilibration for 25 ns and production for 50 ns. The free energy profile (Potential of Mean Force) was then generated with the WHAM program (8) using 90 bins with a convergence tolerance of $10^{-8}$ ensuring overlap between each bin.

*Symmetry-Adapted Perturbation Theory (SAPT)*

In order to analyze non-covalent interactions between the PCB chromophore and the protein environment, Symmetry-Adapted Perturbation Theory (SAPT) (9, 10) was applied to a total of 55 selected snapshots from MD. These structures were obtained from classical MD simulations of SPH and DPH models, sampled between 250 ns to 800 ns in steps of 10 ns. In the SAPT analysis the non-covalent interactions are decomposed into contributions from electrostatics, exchange, induction and dispersion. The calculations were carried for pairwise interaction of a selected residue and the chromophore, which are referred to as monomers. The monomer energies are treated at the Hartree Fock level and the interaction energies are evaluated using a perturbative expansion of the interaction potential. We have used the variant SAPT0 that treats electrostatics and exchange up to first order, while induction and dispersion up to second order. The energies were evaluated with a cc-pVDZ basis set.

*QM/MM Molecular dynamics*

Two structures of the DPH model, one from substate D-$\alpha_f$ and one from substate D-$\beta_f$, extracted from a classical MD trajectory were used as starting points for QM/MM MD simulation. Further, one snapshot of substate D-$\alpha_f$ from the SPH model, was also chosen as a starting point for QM/MM MD simulations. The QM region was chosen to be the full PCB chromophore (including propionates) and sidechains of the three residues in the binding pocket namely C321, D291 and H322. The single bond between $C_\alpha$ and $C_\beta$ atoms was truncated and capped in all residues. Thus, in total, the QM region contained 106 atoms for the DPH model and 105 atoms for the SPH model. The MM region consisted of the remaining protein and the solvent molecules. The QM region was described by DFTB2 with dispersion correction (11, 12) while the MM region was described by AMBER ff14SB force field. Molecular dynamics was carried out for 1 ns with a time step of 1 fs. In the DPH model, an unusual proton transfer from the δ-nitrogen of H322 to one of the oxygen atoms of the carboxylate group in the propionate chain at the B-ring was observed within a short time-scale of 10 ps in the simulations of both substates. However, no such proton transfer was observed in the simulation of substate D-$\alpha_f$ in the SPH model. Snapshots sampled every 10 ps from QM/MM MD were then used to generate input files using the AMBER-ORCA (13) interface to compute absorption and CD spectra. Spectra were calculated on two different QM regions namely: QM106 (DPH model)/QM105(SPH model), which consisted of PCB, C321, H322 and D291 residues and QM66 which consisted of only the four pyrrole rings of the PCB chromophore. For very large organic molecules, sTD-DFT with a hybrid range separated functional is reported to produce accurate absorption and CD spectra in a reasonable amount of computation time (14–16). It has shown to perform quite well for the PCB chromophore (17). Thus, for the sTD-DFT calculations we have used the hybrid range separated functional CAM-B3LYP with a def2-SVP basis set for calculations of excitation energies and rotatory strengths similar to our previous study of Slr1393g3 (18). The excitation energies computed with sTD-DFT as implemented in ORCA version 4.0 were further validated by ADC(2)/cc-pVDZ calculations for both DPH and SPH models with Turbomole 7.2. However, owing to the high computational demand in ADC(2), we calculated excitation energies and rotatory strengths only for the QM66 region. The absorption and CD spectra were generated by convoluting the first 30 transitions from sTD-DFT and 10 transitions from RI-ADC(2) using a Gaussian function with a broadening of 0.15 eV full width at half maximum. The oscillator strengths and rotatory strengths were considered in the dipole-length representation. The spectra of all snapshots were then averaged.

**Figures**

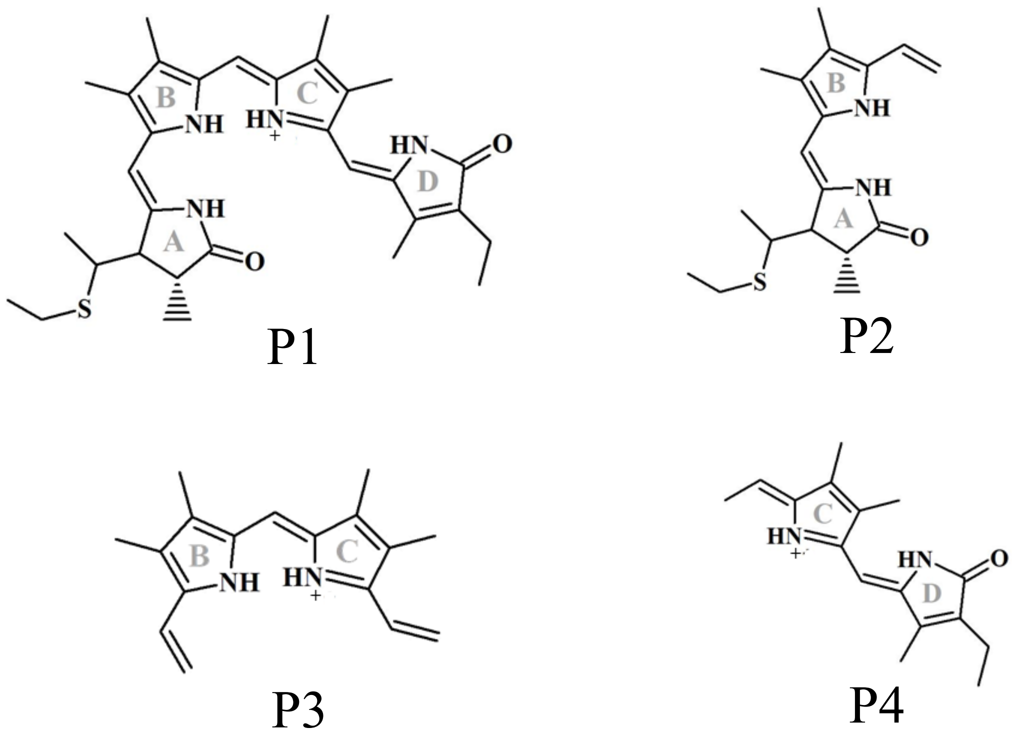


**Fig. S1.** Model Fragments used for parameterization namely: Fragment 1 (P1), Fragment 2 (P2), Fragment 3 (P3) and Fragment 4 (P4).

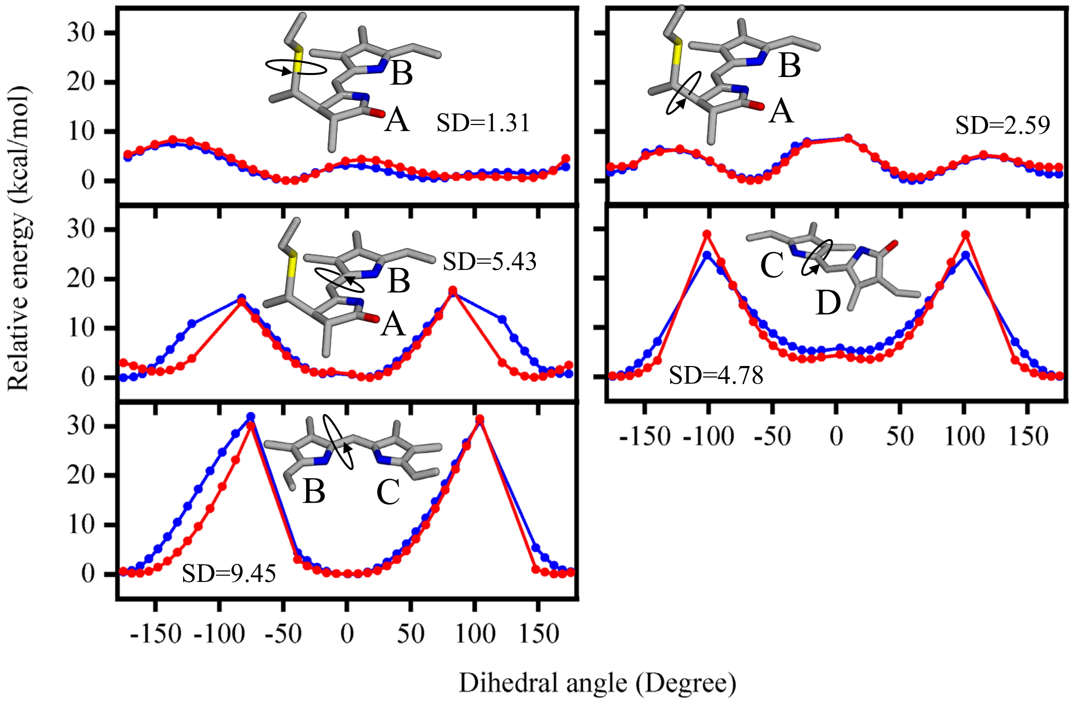


**Fig. S2.** Comparison of QM (red) and MM (blue) profiles in gas-phase. To validate the new developed parameters for the PCB chromophore, MM-minimized conformations of the individual fragments (P2, P3 and P4) were generated by increasing the dihedral angles from -180º to +180º in steps of 10º. All rotational profiles exhibit two minima, except the dihedral between the B and C rings in fragment P3 which shows only one minimum. Single point energies were computed for all conformations using the new force field parameters and the quantum chemical method MP2 with a cc-pVTZ basis set. There is a close agreement between the calculated energy profiles from the parametrized force field (MM) and the MP2 (QM). The parametrization process was focused on the dihedrals around 0° because it is unlikely that an isomerization will take place in the ground state. Hence, we have focused on detailed parametrization around the ground state minimum. The standard deviation (SD) between the QM and the MM profiles is highest for the B-C dihedral with a value of 9.45 kcal/mol and lowest for the dihedral connecting the cysteine and the A-ring with a value of 1.31 kcal/mol considering the region close to the minimum between -100° to +100°.

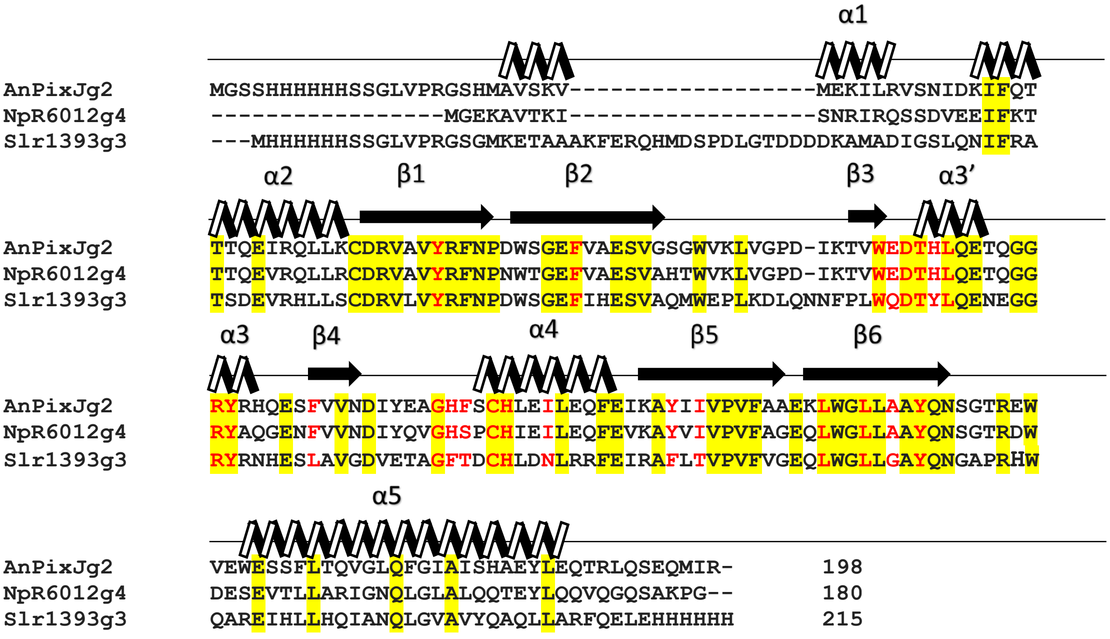


**Fig. S3.** Alignment between the sequences of the red/green CBCRs NpR6012g4, AnPixJg2 and Slr1393g3. Conserved residues between all three are highlighted in yellow. Residues present within 5 Å of the chromophore in AnPixJg2 are highlighted in red.

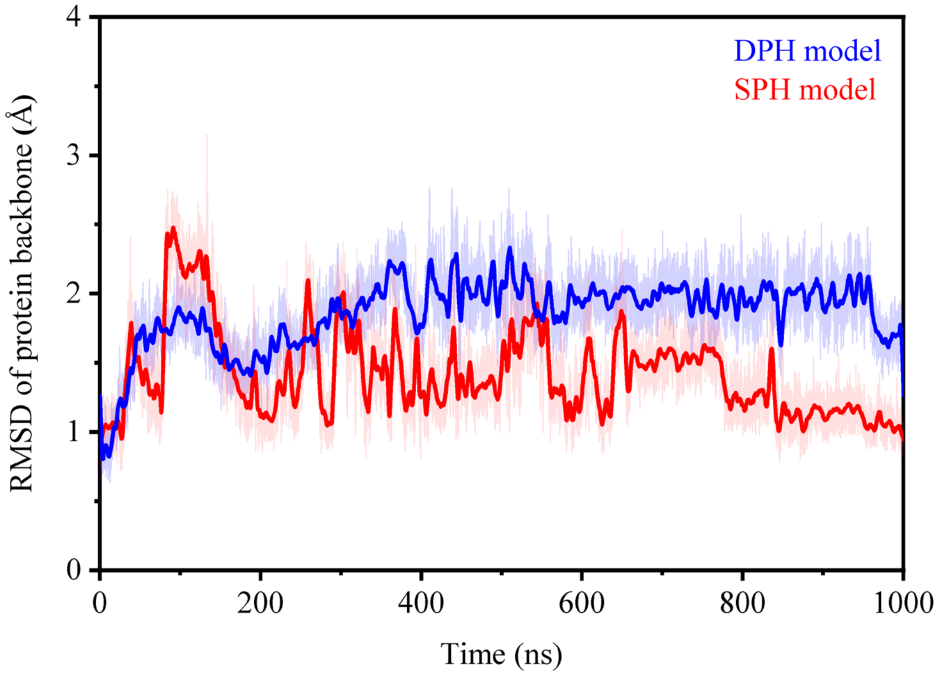


**Fig. S4.** Root-mean-square deviation **(**RMSD) of the protein backbone. The SPH model shows more fluctuations in the geometry during the first 350 ns compared to the DPH model after which the fluctuations reduce thus stabilizing the RMSD. The structural rearrangements of the β2 and β3 sheets is one of the major contributors to RMSD during 300 to 650 ns in both SPH and DPH models.

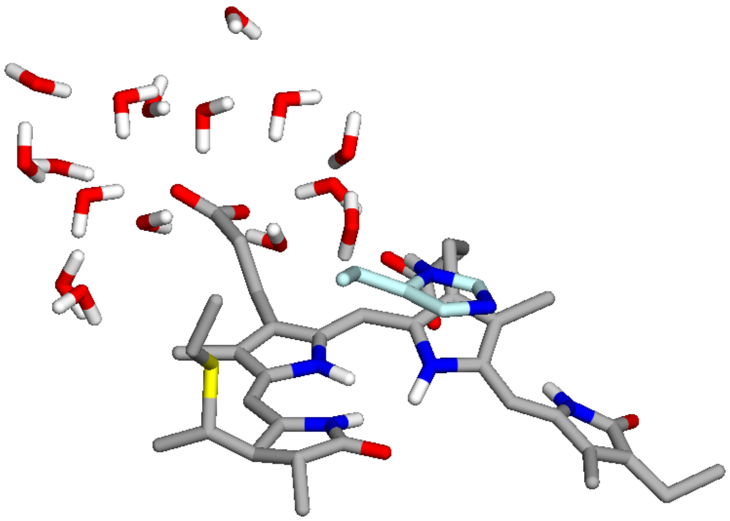

**Fig. S5.** Reorientation of propionate side chain B in the SPH model towards the solvent**.** A Solvent exposed conformation of the B-ring propionate in the D-$\beta_f$ of SPH model was observed during the umbrella sampling contributing to higher PMF.

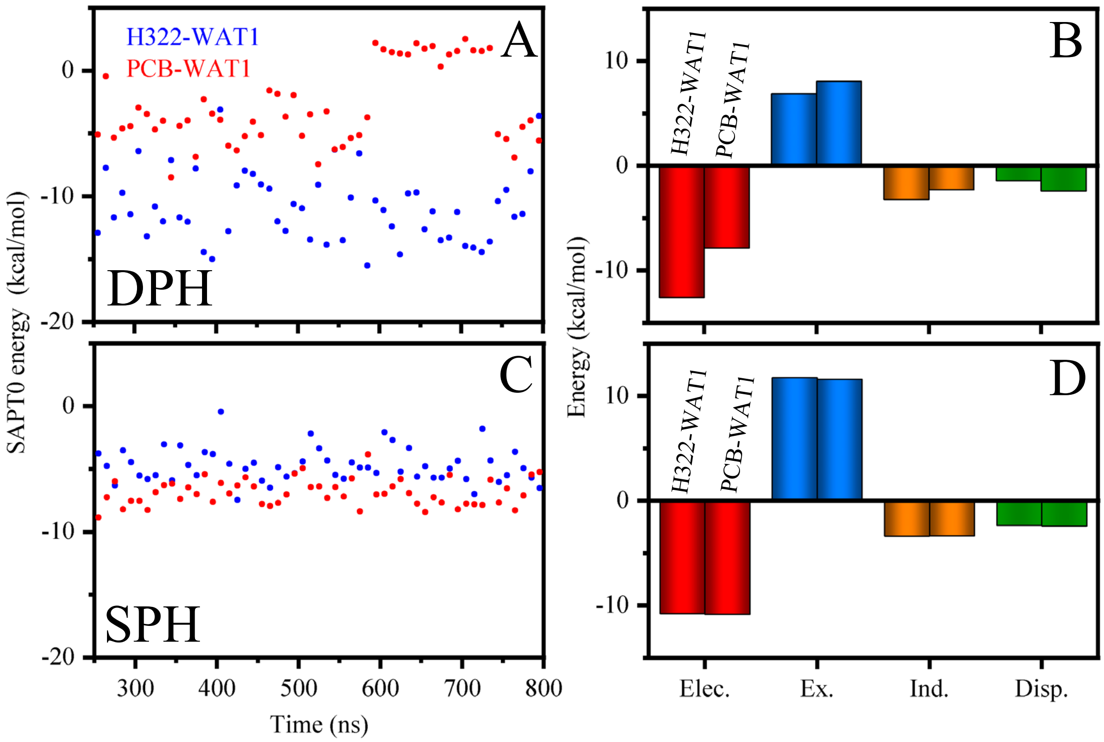


**Fig. S6.** Interaction energies computed for PCB-WAT1 and H322-WAT1. (A) SAPT0 energies of PCB-WAT1 (red) and H322-WAT1 (blue) in the DPH model. The mean SAPT0 energy of H322-WAT1 and that of PCB-WAT1 in substate D-$\alpha_f$ of the DPH model is -10.33 kcal/mol and -4.54 kcal/mol, respectively. (B). SAPT0 energy decomposition of PCB-WAT1 and H322-WAT1 interactions in the DPH model into Electrostatics (red), Exchange (blue), Induction (orange) and Dispersion energies (green). The first column in each energy component in the bar diagram corresponds to the H322-WAT1 interaction and the second column corresponds to PCB-WAT1 interaction. (C). SAPT0 energies of PCB-WAT1 (red) and H322-WAT1(blue) in the SPH model. The mean SAPT0 energy of H322-WAT1 in substate D-$\alpha_f$ of the SPH model is -4.74 kcal/mol and that of PCB-WAT1 is -6.89 kcal/mol. (D). SAPT0 energy decomposition of PCB-WAT1 and H322-WAT1 interactions in the SPH model with the same color coding as in (B).

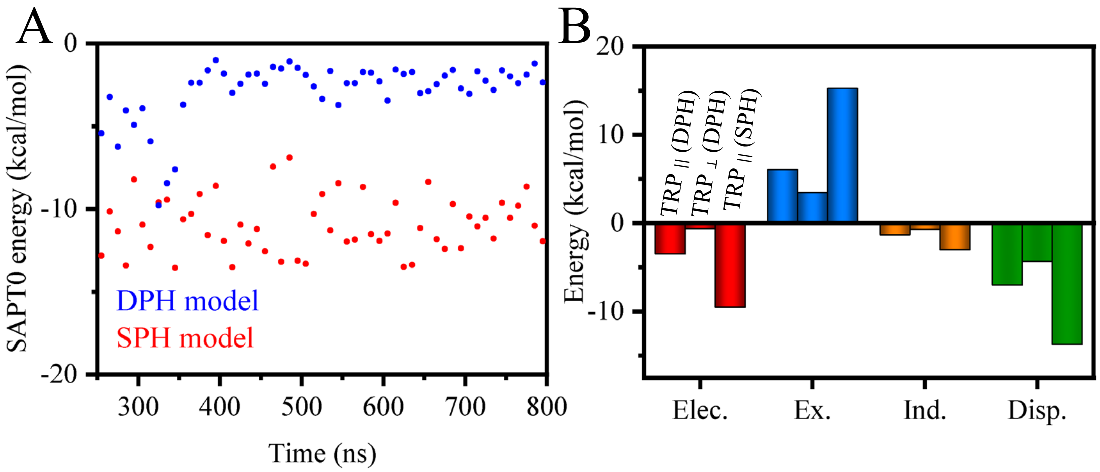


**Fig. S7.** Interaction energies computed for PCB-W289. **(A).** SAPT0 energies between PCB and W289 in DPH (blue) and SPH (red) models. In the DPH model, the mean SAPT0 energy of PCB-W289 with W289 in the parallel conformation is -5.75 kcal/mol and with W289 in the perpendicular conformation is -2.15 kcal/mol. In the SPH model, the value is -10.95 kcal/mol. (B). SAPT0 energy decomposition of PCB-W289 interactions in DPH and SPH models into Electrostatics (red), Exchange (blue), Induction (orange) and Dispersion energies (green). The first column and second columns of the energy components in the bar diagram correspond to the parallel and perpendicular conformations of W289 in the DPH model while the third column corresponds to the parallel conformation of W289 in the SPH model.

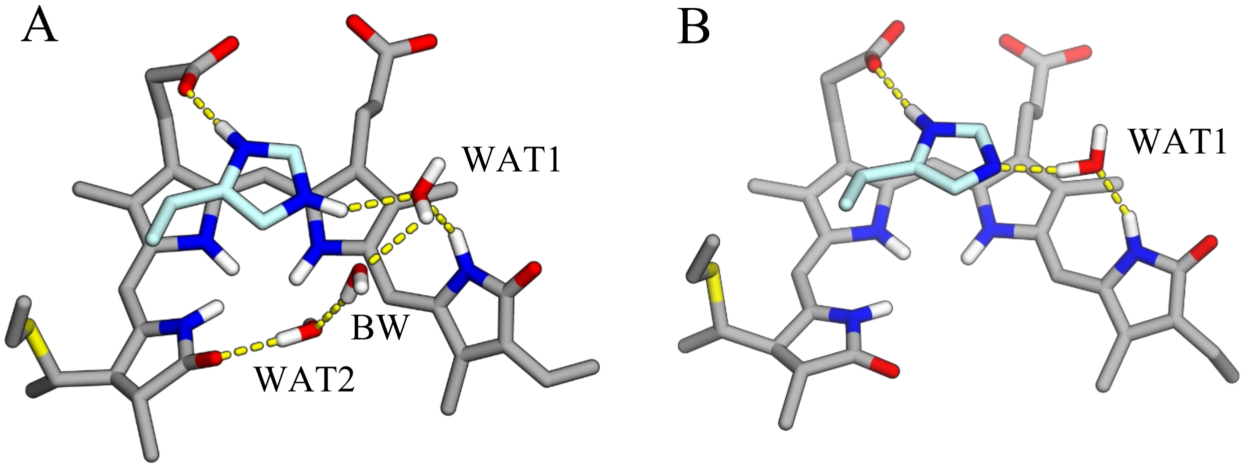


**Fig. S8.** Typical Water networks observed during the simulations of the (A) DPH model and (B) SPH model. In the DPH model, a bridging water (BW) molecule connects the WAT1 and WAT2 molecules. In the SPH model, WAT2 has a very low occurrence and the BW was not observed.

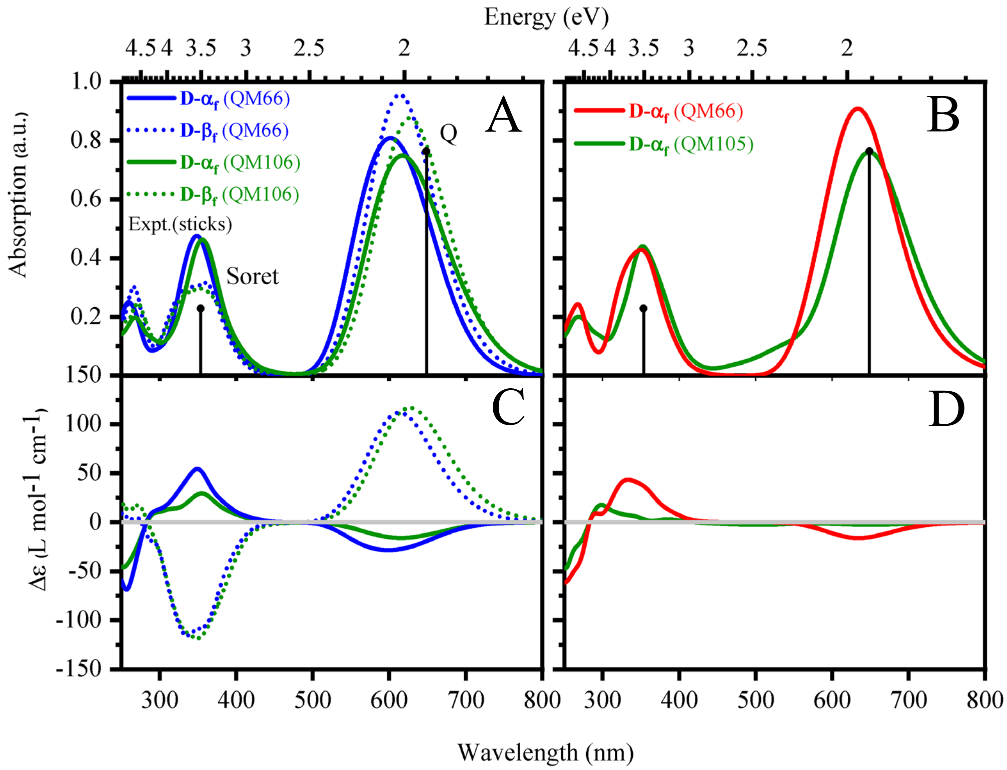


**Fig. S9.** Absorption and CD spectra computed at sTD-DFT/CAM-B3LYP level of theory. Substate D-$\alpha_f$ (solid) and substate D-$\beta_f$ (dashed) of DPH model is shown in A and C, while the spectra of the SPH model are shown in B and D (red or green). The bigger QM region (QM106 (DPH model)/ QM105 (SPH model) is shown in green color in both models. The spectra are averages from 100 snapshots for each graph.

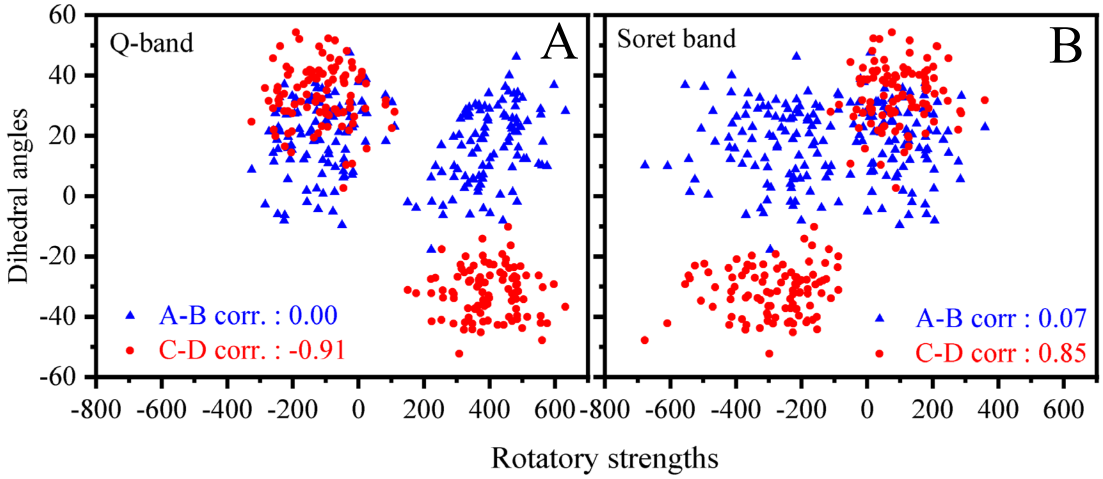


**Fig. S10.** Correlation of the A-B (black) and C-D (red) dihedral angles over the rotatory strengths computed from ADC(2)/cc-pVDZ for both substate D-$\alpha_f$ and D-$\beta_f$ of the DPH model. (A). Rotatory strengths comprising the Q-band which is mainly the $S_0 \rightarrow S_1$ transition. (B). Rotatory strengths comprising the Soret-band which comprises of summed up contributions from $S_0 \rightarrow S_2$, $S_0 \rightarrow S_3$, $S_0 \rightarrow S_4$ and $S_0 \rightarrow S_5$ transitions.

### Tables

**Table S1.** Mean and standard deviation of dihedrals constituting the methine bridges in SPH and DPH models

| **Dihedrals around the methine bridge** | **Crystal structure values** | **DPH model** | | | | **SPH model** | |
|---|---|---|---|---|---|---|---|
| | | **Substate D-$\alpha_f$** | | **Substate D-$\beta_f$** | | **Substate D-$\alpha_f$** | |
| | | **Mean** | **Std. dev.** | **Mean** | **Std. dev.** | **Mean** | **Std. dev.** |
| A-B single | 10.58 | 18.55 | 9.12 | 19.28 | 8.60 | 12.42 | 10.35 |
| A-B double | 0.38 | 3.26 | 6.86 | 3.57 | 6.76 | 1.89 | 6.83 |
| B-C single | 2.21 | 2.95 | 6.34 | 1.30 | 6.30 | 4.72 | 6.28 |
| B-C double | -6.58 | 7.19 | 6.82 | 4.52 | 6.83 | 8.18 | 6.81 |
| C-D single | 36.08 | 31.13 | 9.97 | -23.63 | 9.18 | 30.58 | 9.46 |
| C-D double | 26.96 | 14.26 | 7.77 | -8.39 | 7.20 | 13.13 | 7.60 |

**Table S2.** Hydrogen bonding occurrences in SPH and DPH models

| Residue | Donor | Acceptor | DPH model | | SPH model |
|---|---|---|---|---|---|
| | | | **Sub-state I** | **Sub-state II** | **Sub-state I** |
| D291 | NC | OD1 | 0.51 | 0.49 | 0.22 |
| | | OD2 | 0.00 | 0.00 | 0.38 |
| | ND | OD1 | 0.53 | 0.82 | 0.13 |
| | | OD2 | 0.00 | 0.00 | 0.34 |
| | NA | OD1 | 0.99 | 0.99 | 0.30 |
| | | OD2 | 0.00 | 0.00 | 0.66 |
| W289 | NE1 | OC | 0.05 | 0.00 | 0.31 |
| H322 | ND1 | O1D | 0.55 | 0.54 | 0.44 |
| | | O2D | 0.52 | 0.47 | 0.41 |
| H318 | ND1 | O1D | 0.25 | 0.33 | 0.40 |
| | | O2D | 0.25 | 0.40 | 0.41 |
| Y352 | OH | OB | 0.56 | 0.00 | 0.83 |
| R301 | NH2 | O1A | 0.54 | 0.37 | 0.42 |
| | NH2 | O2A | 0.47 | 0.42 | 0.61 |
| | NE | O1A | 0.40 | 0.21 | 0.56 |
| | NE | O2A | 0.48 | 0.36 | 0.32 |
| Y302 | OH | OB | 0.00 | 0.46 | 0.00 |
| | | O1A | 0.37 | 0.00 | 0.56 |
| | | O2A | 0.44 | 0.00 | 0.32 |
| Water-D ring | NB | O | 0.58 | 0.66 | 0.85 |
| Water-A ring | O | OC | 0.79 | 0.72 | 0.06 |

**Table S3.** List of atoms, atom types and partial charges. Atoms of cysteine moiety are assigned AMBER atom type and those of PCB are assigned GAFF type.

| Atom name | Atom type | Partial charge | Atom name | Atom type | Partial charge |
|---|---|---|---|---|---|
| N | N | -0.4157 | C2B | cd | -0.06711 |
| H | H | 0.2719 | CMB | c3 | -0.22662 |
| CA | CX | 0.0743 | HMB1 | hc | 0.10421 |
| CB | 2C | 0.02777 | HMB3 | hc | 0.10421 |
| HB3 | H1 | 0.05705 | HMB2 | hc | 0.10421 |
| HB2 | H1 | 0.05705 | CAB | c3 | 0.01705 |
| SG | S | -0.25459 | HAB1 | hc | 0.05755 |
| CAC | c3 | 0.04195 | HAB2 | hc | 0.05755 |
| C3C | c3 | -0.16391 | CBB | c3 | -0.13539 |
| C2C | c3 | 0.12596 | HBB3 | hc | 0.05367 |
| C1C | c | 0.45191 | HBB2 | hc | 0.05367 |
| NC | na | -0.47493 | HBB1 | hc | 0.05367 |
| C4C | ca | 0.26663 | OB | o | -0.57233 |
| CHD | m1 | -0.3885 | HB | hn | 0.45348 |
| HHD | ha | 0.19106 | HHA | ha | 0.14555 |
| C1D | cc | 0.20004 | C3D | cd | 0.19796 |
| ND | na | -0.41051 | C2D | cd | -0.04998 |
| HD | hn | 0.34477 | CMD | c3 | -0.19296 |
| C4D | cc | -0.07335 | HMD2 | hc | 0.07943 |
| CHA | m2 | 0.01904 | HMD3 | hc | 0.07943 |
| C1A | cd | 0.01388 | HMD1 | hc | 0.07943 |
| NA | na | -0.52954 | CAD | c3 | -0.19321 |
| H70 | hn | 0.38913 | CBD | c3 | -0.1722 |
| C4A | cd | 0.32176 | HBD2 | hc | -0.0122 |
| C3A | cc | -0.12451 | CGD | c | 0.7994 |
| C2A | cc | 0.10876 | O2D | o | -0.8014 |
| CAA | c3 | -0.20179 | O1D | o | -0.8014 |
| CBA | c3 | -0.1722 | HBD1 | hc | -0.0122 |
| HBA2 | hc | -0.0122 | HAD1 | hc | 0.11639 |

| | | | | | |
|---|---|---|---|---|---|
| CGA | c | 0.7994 | HAD2 | hc | 0.11639 |
| O2A | o | -0.8014 | HC | hn | 0.39155 |
| O1A | o | -0.8014 | OC | o | -0.5105 |
| HBA1 | hc | -0.0122 | H2C | hc | 0.06038 |
| HAA1 | hc | 0.11963 | CMC | c3 | -0.26137 |
| HAA2 | hc | 0.11963 | HMC3 | hc | 0.08866 |
| CMA | c3 | -0.17282 | HMC2 | hc | 0.08866 |
| HMA3 | hc | 0.08269 | HMC1 | hc | 0.08866 |
| HMA1 | hc | 0.08269 | H3C | hc | 0.11156 |
| HMA2 | hc | 0.08269 | HAC2 | h1 | 0.11809 |
| CHB | m3 | -0.50431 | CBC | c3 | -0.26952 |
| HHB | ha | 0.2278 | HBC2 | hc | 0.09265 |
| C1B | m4 | 0.46312 | HBC3 | hc | 0.09265 |
| NB | na | -0.78804 | HBC1 | hc | 0.09265 |
| C4B | c | 0.82991 | HA | H1 | 0.0766 |
| C3B | cd | -0.19499 | C | C | 0.5973 |
| | | | O | O | -0.5679 |

**Table S4.** Optimized parameters for angles

| **Angle Parameter** | **Force constant for angle** $(k_w)$ | **Equilibrium value** $(\theta_0)$ |
|---|---|---|
| ha-m1-cc | 47.2766 | 120.52 |
| ca-m1-ha | 42.5876 | 118.76 |
| ca-m1-cc | 66.6056 | 118.32 |
| ha-m2-cd | 45.9972 | 119.38 |
| ha-m2-cc | 48.8458 | 121.63 |
| m3-m4-cd | 69.5530 | 122.71 |
| ha-m3-m4 | 43.3257 | 120.91 |
| cd-m3-m4 | 67.0643 | 119.57 |
| cc-m2-cd | 66.3973 | 120.26 |

**Table S5.** Optimized parameters for dihedrals

| Dihedral parameter | Dihedral force constant ($k_j$) | Phase shift ($\gamma$) | Periodicity ($j$) |
|---|---|---|---|
| 2C-S-C3-H1 | 0.86712 | 0 | 3 |
| 2C-S-C3-C3 | 0.00048 | 0 | 3 |
| na-ca-m1-ha | 10.92344 | 180 | 2 |
| na-ca-m1-cc | 8.44994 | 180 | 2 |
| c3-ca-m1-ha | 3.90155 | 180 | 2 |
| c3-ca-m1-cc | 9.23158 | 180 | 2 |
| ha-m1-cc-na | 5.06283 | 180 | 2 |
| ha-m1-cc-cd | 1.79702 | 180 | 2 |
| ca-m1-cc-na | 5.04017 | 180 | 2 |
| ca-m1-cc-cd | 3.79821 | 180 | 2 |
| ha-m2-cc-na | 5.47416 | 180 | 2 |
| ha-m2-cc-cd | 5.92299 | 180 | 2 |
| ha-m2-cd-na | 7.86036 | 180 | 2 |
| ha-m2-cd-cc | 3.77629 | 180 | 2 |
| cd-cc-m2-cd | 3.38927 | 180 | 2 |
| na-cc-m2-cd | 10.27961 | 180 | 2 |
| cc-m2-cd-na | 6.51739 | 180 | 2 |
| cc-m2-cd-cc | 2.2914 | 180 | 2 |
| na-cd-m3-ha | 4.16984 | 180 | 2 |
| na-cd-m3-m4 | 6.64016 | 180 | 2 |
| cc-cd-m3-m4 | 4.31623 | 180 | 2 |
| cc-cd-m3-ha | 4.04927 | 180 | 2 |
| ha-m3-m4-cd | 2.68604 | 180 | 2 |
| ha-m3-m4-na | 8.93888 | 180 | 2 |
| cd-m3-m4-na | 5.57597 | 180 | 2 |
| cd-m3-m4-cd | 4.63996 | 180 | 2 |
